# Single-nucleus RNA-seq of normal-appearing brain regions in relapsing-remitting vs. secondary progressive multiple sclerosis

**DOI:** 10.1101/2022.01.10.475705

**Authors:** Yasuyuki Kihara, Yunjiao Zhu, Deepa Jonnalagadda, William Romanow, Carter Palmer, Benjamin Siddoway, Richard Rivera, Ranjan Dutta, Bruce D. Trapp, Jerold Chun

**Affiliations:** Translational Neuroscience Initiative, Sanford Burnham Prebys Medical Discovery Institute, La Jolla, CA 92037, USA; Department of Neuroscience, Lerner Research Institute, Cleveland Clinic, Cleveland, OH 44195, USA; Biomedical Sciences Graduate Program, School of Medicine, University of California San Diego, 9500 Gilman Dr, La Jolla, CA 92093, USA; Illumina, San Diego, CA 92121, USA; Vertex Pharmaceuticals, San Diego, CA 92121, USA

**Keywords:** Neuroinflammation, S1P_1_, FTY720, siponimod, ozanimod, ponesimod, lysophospholipid receptors

## Abstract

Multiple sclerosis (MS) is an immune-mediated demyelinating disease that alters central nervous system (CNS) functions. Relapsing-remitting MS (RRMS) is the most common form, which can transform into secondary-progressive MS (SPMS) that is associated with progressive neurodegeneration. Single-nucleus RNA sequencing (snRNA-seq) of MS lesions identified disease-related transcriptomic alterations; however, their relationship to non-lesioned MS brain regions has not been reported and which could identify prodromal or other disease susceptibility signatures. Here, snRNA-seq was used to generate high-quality RRMS vs. SPMS datasets of 33,197 nuclei from 8 normal-appearing MS brains, which revealed divergent cell type-specific changes. Notably, SPMS brains downregulated astrocytic sphingosine kinases (*SPHK1/2*) – the enzymes required to phosphorylate and activate the MS drug, fingolimod. This reduction was modeled with astrocyte-specific *Sphk1/2* null mice in which fingolimod lost activity, supporting functionality of observed transcriptomic changes. These data provide an initial resource for studies of single cells from non-lesioned RRMS and SPMS brains.

## Introduction

Single cell transcriptomics has become a principal gateway to understand cellular states and characteristics in normal and diseased tissues (Lahnemann et al., 2020; Regev et al., 2017). Approaches for analyzing single brain cells by single nucleus RNA-sequencing (snRNA-seq) (Fan et al., 2016; Gole et al., 2013; Lake et al., 2016; Lake et al., 2018; Lake et al., 2017; Zheng et al., 2017) have successfully generated reference datasets from non-diseased and diseased human brains that include Alzheimer’s disease (Mathys et al., 2019), Parkinson’s disease (Aneichyk et al., 2018), amyotrophic lateral sclerosis (Maniatis et al., 2019) and multiple sclerosis (MS) (Beltran et al., 2019; Jakel et al., 2019; Masuda et al., 2019; Schirmer et al., 2019).

MS is a neuroinflammatory disease that produces neurodegeneration. It is characterized pathologically by inflammation and demyelination that produce white matter plaque lesions and neuronal damage/loss in the CNS (Hickey, 1999; Thompson et al., 2018). Relapsing-remitting MS (RRMS) is the most common disease course, which can transition into secondary progressive MS (SPMS) that is characterized by progressive neurological disability (Hickey, 1999; Thompson et al., 2018). Pathophysiological differences have been reported between RRMS vs. progressive forms of MS, contrasting with subtypes of progressive MS – primary progressive MS (PPMS) and SPMS – that share similar neuropathologies (Lassmann, 2018).

MS lesions from patient samples that have been analyzed by snRNA-seq thus far have generally been compared to non-diseased brains. Reported features of RRMS include loss of excitatory neurons in the upper-cortical layers, and gene expression profiles characteristic of stressed oligodendrocytes, reactive astrocytes, and activated microglia (Schirmer et al., 2019). In SPMS lesions, there is oligodendroglial heterogeneity, with reduced numbers of oligodendrocyte precursor cells (OPCs) and loss of *OPALIN^+^* oligodendrocyte sub-populations (Jakel et al., 2019). Another study that compared nuclei isolated from tumefactive MS lesions highlighted differences between activated microglia vs. healthy microglia of non-diseased controls (Masuda et al., 2019). These snRNA-seq studies comparing changes between MS brain lesions vs. control brains focused preferentially on cells within and around lesions that lacked comparable cellular controls, as indicated by markedly different cell populations revealed by tSNE (t-distributed stochastic neighbor embedding (Jamieson et al., 2010)) or UMAP (uniform manifold approximation and projection (Becht et al., 2018)) clustering plots.

The possibility that there are disease-related transcriptomic changes in the MS brain apart from demyelinating lesions is predicted by the known discordance between lesion burden and clinical presentation, as well as grey matter changes (Geurts et al., 2012) and brain volume loss in MS (Bermel and Bakshi, 2006). To identify possible global changes in disease-validated RRMS and SPMS brains, neuroanatomically-matched, normal-appearing prefrontal cortices with no apparent MS lesions were assessed by snRNA-seq.

## Results

### snRNA-seq identified major CNS cell types from non-lesioned MS brains

A previously developed snRNA-seq pipeline for human postmortem brains was used to assess gene expression signatures in normal-appearing MS brains. None of the 10 collected MS prefrontal cortices (Brodmann area 10; 5 from RRMS and 5 from SPMS) showed detectable demyelination (**Fig. 1A**). No statistical differences in age, sex, or postmortem interval (PMI) were identified between RRMS vs. SPMS. The median expanded disability status scale (EDSS) was higher in SPMS than in RRMS (median EDSS = 9.0 vs. 4.0, respectively: **Supplemental Table S1**). Brain samples with an RNA integrity number (RIN) ≥ 5 (median RIN = 7.35: **Supplemental Table S1**) were sectioned at ~300 μm thickness and processed for fluorescence-activated nuclear sorting (FANS). DAPI^+^ singlet nuclei were barcoded using a 10x Genomics Chromium single cell 3’ reagent kit, followed by short-read sequencing on an Illumina HiSeq3000 instrument (**Fig. 1B**). Quality control filtering resulted in a total of 33,197 nuclei (12,431 nuclei from 3 RRMS brains vs. 20,766 nuclei from 5 SPMS brains) with ~6-fold more transcripts per nucleus (5,977 ± 735 vs. 5,420 ± 725 transcripts/nucleus in RRMS vs. SPMS, respectively; **Supplemental Table S2**) than prior reports (1,096 transcripts/nucleus (Jakel et al., 2019) and 2,400 transcripts/nucleus (Schirmer et al., 2019)). Cells were clustered using the Seurat SNN (shared nearest neighbor) algorithm and visualized as UMAP plots (**Fig. 1C**).

**Figure 1.**
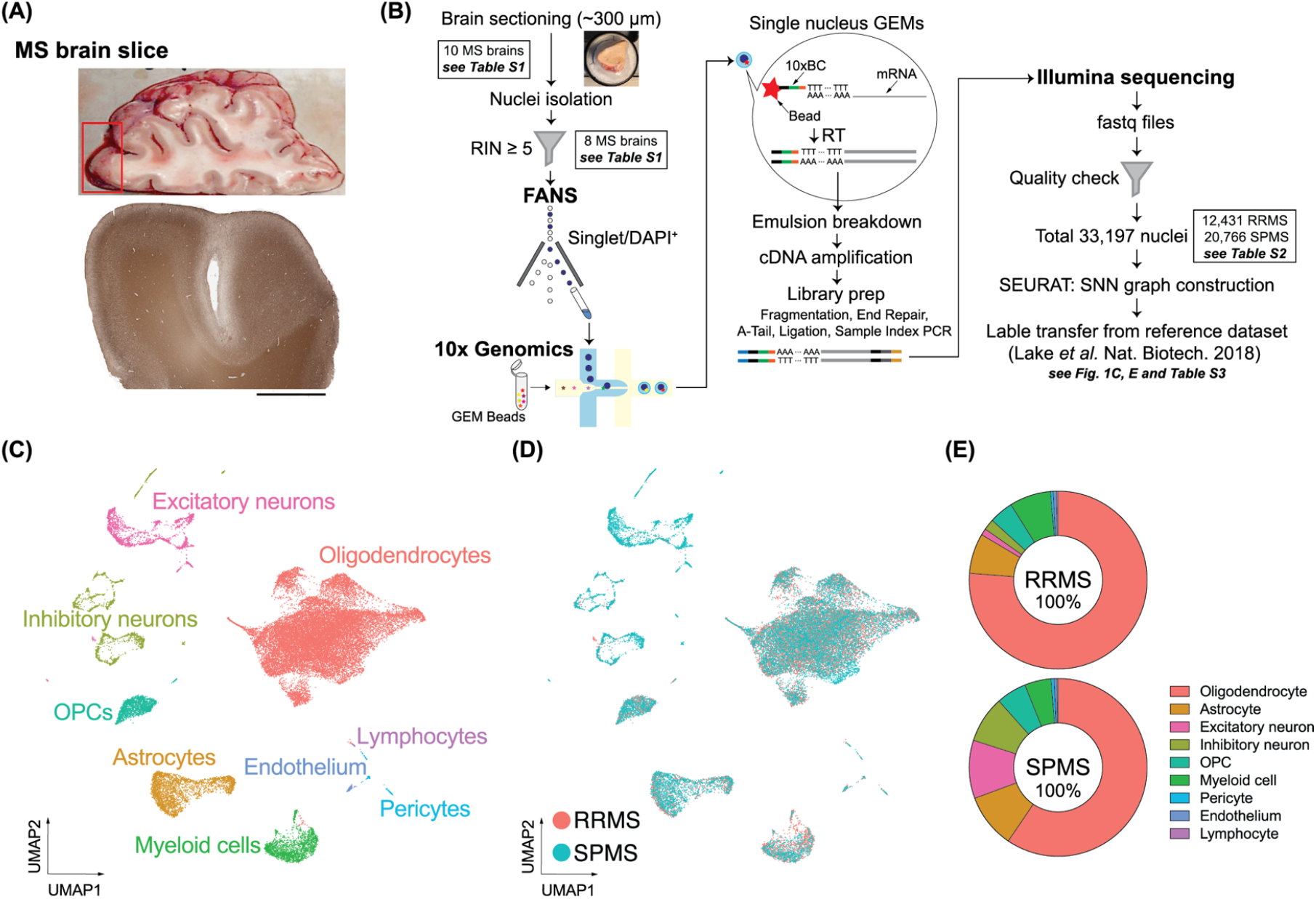
Experimental approach of snRNA-seq and unbiased cellular identification in MS brains. (**A**) Macroscopic and immunohistochemical view of a representative MS brain. (**B**) Experimental workflow. Nuclei were isolated from ~300 μm frozen MS brain sections by fluorescence-activated nuclear sorting (FANS) based on DAPI (4’,6-diamidino-2-phenylindole) positivity. Brain samples with a RNA integrity number (RIN) ≥ 5 were processed for 10X Genomics and snRNA-seq. (**C**) Unbiased cellular identification. Quality-control filtering identified a total of 33,197 nuclei from both RRMS and SPMS brains; these cells were then clustered using the Shared Nearest Neighbor (SNN) clustering algorithm in the SEURAT package. The cell types were annotated based on the reference dataset of normal human brains (Lake et al., 2018) and are presented as a UMAP (Uniform Manifold Approximation and Projection) plot. Colors indicate specific cell clusters identified by SEURAT V.3.0. (**D**) UMAP plots for RRMS and SPMS cells. (**E**) Relative proportions of the 9 cell types analyzed in the RRMS and SPMS brains (see details in **Supplementary Table S3**).

Cell types were determined by label transfer from a reference dataset (Lake et al., 2018), resulting in 9 major cell types including astrocytes, endothelial cells, excitatory and inhibitory neurons, oligodendrocytes (OLs), oligodendrocyte progenitor cells (OPCs), lymphocytes, myeloid cells (microglia/macrophages), and pericytes (**Fig. 1C**). The reference dataset enabled cell type annotation but it was not used for gene expression comparison to avoid uncertain outcomes derived from different experimental techniques and conditions. Cell types were confirmed by the expression patterns of well-established marker genes (**Supplemental Fig. S1**). UMAP plots showed consistent and largely overlapping layouts between RRMS vs. SPMS (**Fig. 1D**, **Fig. 1E and Supplemental Table S3**). Neurons (RRMS ~3%; SPMS ~17% of total) were less represented than in prior non-diseased studies (~75% neurons) (Lake et al., 2018; Schirmer et al., 2019) because neuropathological sample isolation targeted cerebral cortical grey matter-to-white matter projections that would typically contain MS plaques. Consistently, most identified cells were OLs in both RRMS and SPMS samples (~70% of total). Lymphocyte clusters in normal-appearing MS brains (~0.20% of total) were generally comparable to those in prior reports (~0.6% in MS brains as compared to 0% in control non-diseased brains (Schirmer et al., 2019)).

### Excitatory neurons display increased vulnerability in RRMS over SPMS

Comparisons of gene expression profiles between the RRMS vs. SPMS excitatory neurons identified 1,851 differentially expressed genes (DEGs) (**Fig. 2A and Supplemental Table S4**). Marker genes for excitatory neurons (*CBLN2, GLIS3, CUX2, RORB, IL1RAPL2, TSHZ2, FOXP2, PCP4, HS3ST2*) were significantly downregulated in RRMS compared to SPMS (**Fig. 2B**). In addition to DEGs, cell detection rates (CDRs) were calculated as the proportion of cells that express particular genes within each cell cluster (**Supplemental Table S5**). The CDRs in combination with DEGs enabled the identification of vastly altered genes (VAGs) that represented alterations in both expression level and frequency (**Supplemental Table S6**). These analyses identified 837 VAGs in excitatory neurons that included upper layer marker genes (*CUX2, CBLN2, RBFOX3, SATB2*) whose CDRs were decreased in RRMS as compared to SPMS (**Fig. 2C and Supplemental Table S6**). Reactome pathway analyses of upregulated DEGs in RRMS revealed enrichment of the immune response-related pathways (*e.g*., antigen presentation, interferon pathways: **Supplemental Table S7**). The neuronal vulnerability of RRMS brains, which might reflect prior loss of susceptible RRMS neurons in SPMS, may produce neural circuit rewiring issues during disease progression, resulting in cognitive impairment in SPMS (Benedict et al., 2020; Brochet and Ruet, 2019).

**Figure 2.**
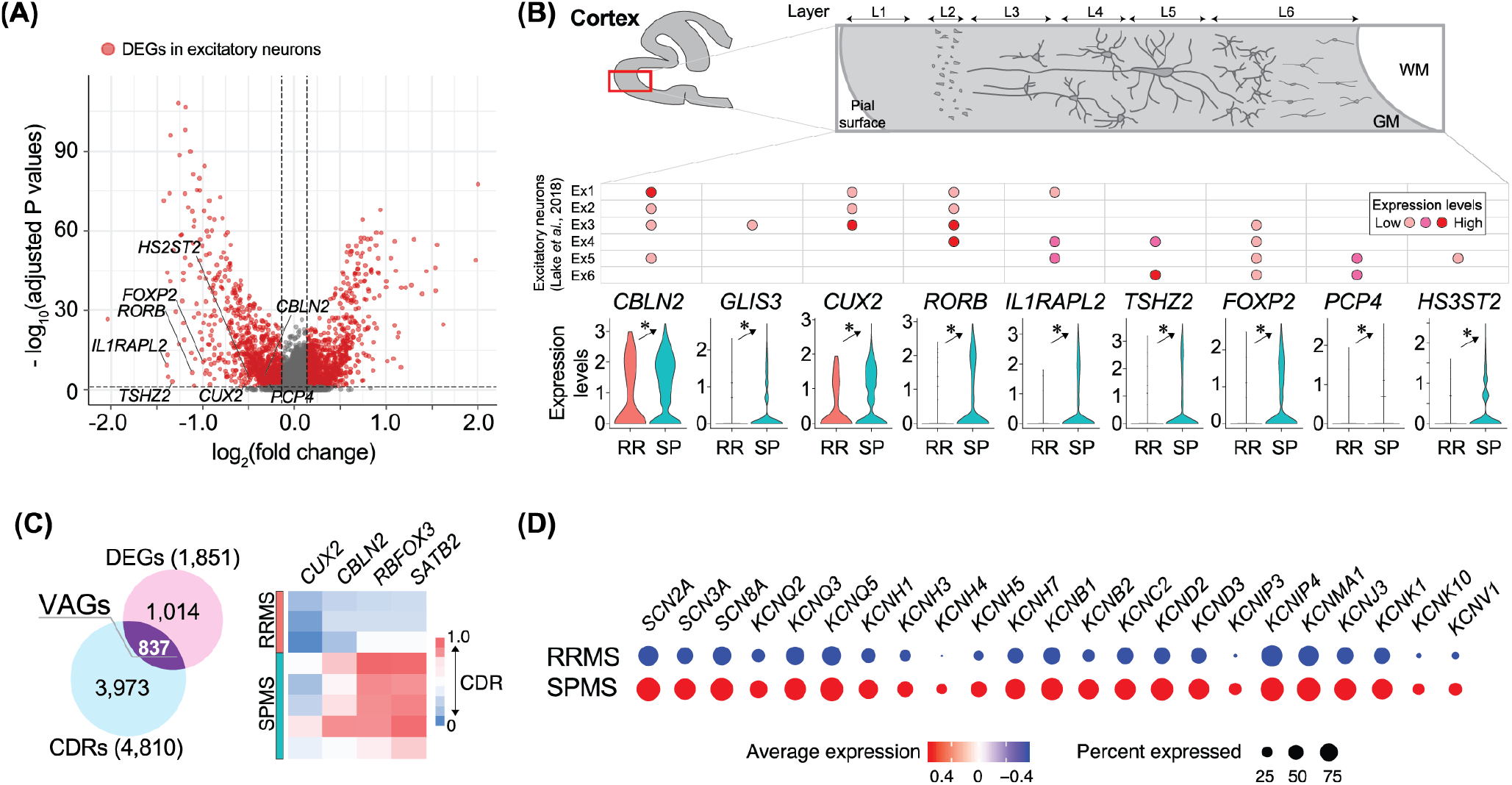
Transcriptomic divergence in neuronal populations between RRMS vs. SPMS. (**A**) Volcano plots of genes identified in excitatory neurons. *Red dots* represent differentially expressed genes (DEGs; adjusted p values < 0.05 and fold change > 1.1) and *grey dots* represent non-significant genes. (**B**) Violin plots of excitatory neuron marker gene expression levels and a table for their expression patterns in the cortical layer determined by previous study (Lake et al., 2018). *, adjusted p value < 0.05. (**C**) Venn diagram of DEGs and genes of differential CDRs (cell detection rates; the proportion of cells that express particular genes within each cell cluster), and a heatmap of excitatory neuron marker genes found in VAGs (vastly altered genes). (**D**) Dot plots of expression levels of sodium and potassium channel genes. Size and color indicate CDRs and expression levels, respectively. Statistical data for DEGs, CDRs and VAGs are provided in **Supplemental Table**.

In SPMS, excitatory neurons showed enrichment of synaptic transmission pathways (*e.g*., neurexins/neuroligins, channels, axon guidance: **Supplemental Table S7**). Notably, DEGs associated with sodium and potassium channels (**Fig. 2D**) were highly upregulated in SPMS, which might reflect potential channelopathy-like defects in SPMS (Kumar et al., 2016; Roostaei et al., 2016; Zhong et al., 2016) because impairment of channels causes axonal degeneration (Dutta and Trapp, 2014).

### OLs show decreased vulnerability and increased stress response genes in RRMS over SPMS

OLs were characterized by the expression of *PLP1, MOG*, and *MAG* (**Fig. 3A, B and Supplemental Fig. S1**). These genes, as well as *RTN4* (encoding Nogo-A) and *S1PR5* (**Fig. 3A and B**), were significantly higher in RRMS vs. SPMS. OL depletion was not observed in these normal-appearing samples, contrasting with their loss from MS lesions (Jakel et al., 2019), while lower expression levels of OL marker genes in SPMS suggested a potential vulnerability of OLs (Dutta and Trapp, 2014), which may lead to OL loss in and around developing active SPMS lesions.

**Figure 3.**
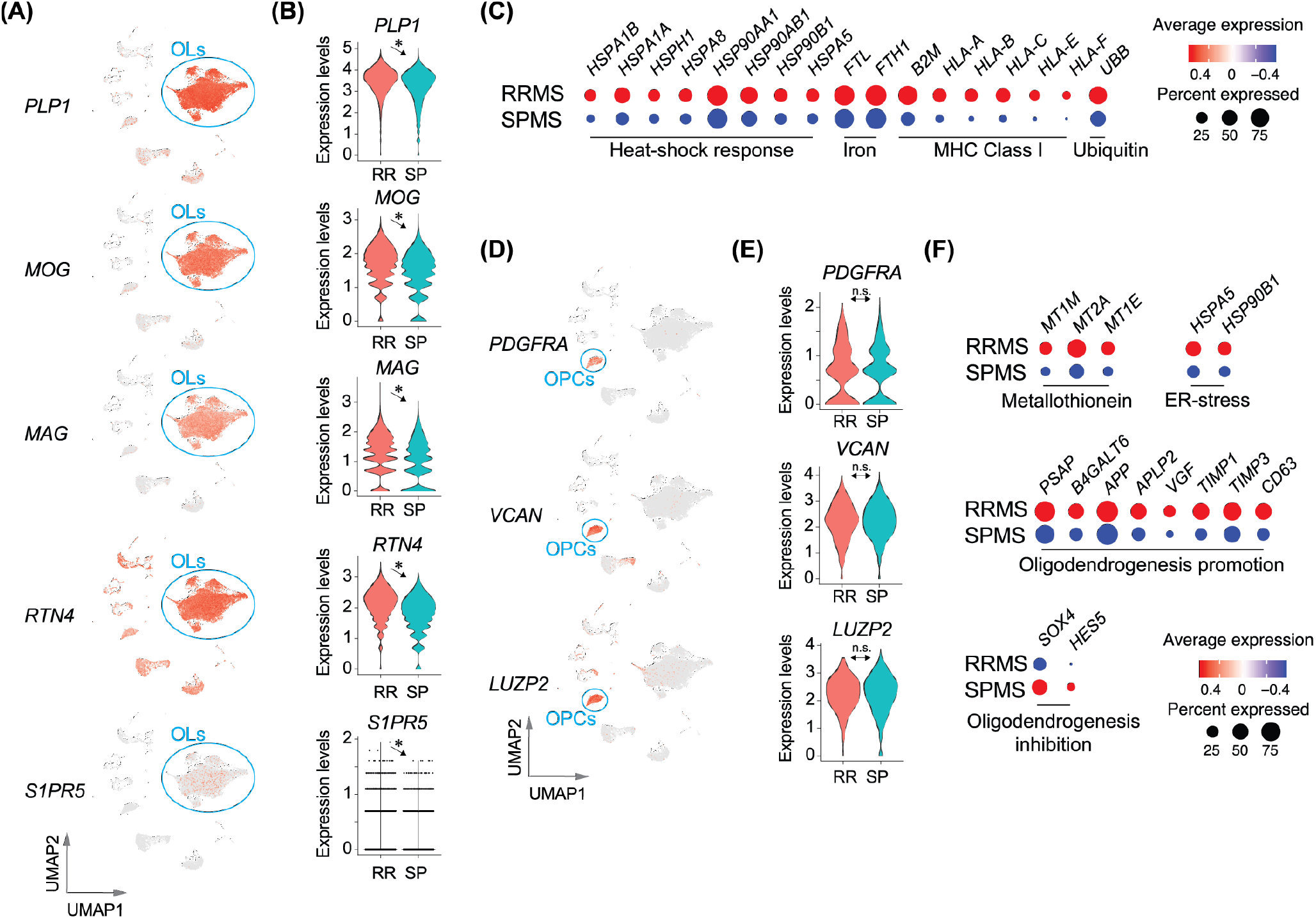
Transcriptomic pathological signatures of MS oligodendrocytes (OLs) and oligodendrocyte progenitor cells (OPCs). (**A and B**) UMAP (A) and violin plots (B) of OL marker genes. *, adjusted p value < 0.05. (**C**) Dot plots for genes associated with heat-shock response, iron accumulation, major histocompatibility complex class I, and ubiquitin-mediated protein degradation. Size and color indicate the CDRs and expression levels, respectively. (**D and E**) UMAP (D) and violin plots (E) of OPC marker genes. n.s., non-significant. (**F**) Dot plots of genes associated with metallothionein pathway genes, ER-stress related genes, and oligodendrogenesis-promoting or inhibiting genes. Size and color indicate frequency and expression levels, respectively. Statistical results for DEGs, CDRs and VAGs are provided in **Supplemental Table**.

OLs showed the third-highest number of DEGs (1,145), ~94% of which were markedly upregulated in RRMS vs. SPMS (**Supplemental Table S4**) and included genes associated with heat-shock response (*HSPA1B, HSPA1A, HSPH1, HSPA8, HSP90AA1, HSP90AB1, HSP90B1*, and *HSPA5*), iron accumulation (*FTL* and *FTH1*), major histocompatibility complex class I (*B2M, HLA-A, HLA-B, HLA-C, HLA-E*, and *HLA-F*), and ubiquitin-mediated protein degradation (*UBB*) (**Fig. 3C**). Reactome pathway analyses of the extracted 158 VAGs (**Supplemental Table. S5 and S6**) highlighted the nonsense-mediated mRNA decay (NMD) pathway that functions in RNA surveillance to degrade aberrant mRNAs (**Supplemental Table S8**). The expression of NMD antagonist *UPF3A* in RRMS OLs was elevated as compared to SPMS, indicating an accumulation of aberrant RNAs produced by low NMD in RRMS OLs (Shum et al., 2016). Collectively, RRMS OLs might reflect more severe cellular stress, leading to subsequent OL vulnerability in SPMS.

### OPCs express more maturation genes in RRMS over SPMS

OPCs (RRMS 4.6%; SPMS 5.8% of total) characterized by expression of *PDGFRA, VCAN*, and *LUZP2* (**Fig. 3D, E and Supplemental Fig. S1**) presented 243 DEGs (**Supplemental Table S4**), 34 of which were extracted as VAGs (**Supplemental Fig. S2**). Reactome pathway analyses identified upregulation of metallothionein pathway genes (*MT1M, MT2A*, and *MT1E*) and ER-stress related genes (*HSPA5* and *HSP90B1*) in RRMS (**Fig. 3F and Supplemental Table S9**). Importantly, elevation of myelin formation-promoting genes (*PSAP, B4GALT6, APP, APLP2, VGF, TIMP1, TIMP3, CD63*) (Alvarez-Saavedra et al., 2016; Hiraiwa et al., 1999; Nicaise et al., 2019; Truong et al., 2019; Yoshihara et al., 2018) were accompanied by decreases in oligodendrogenesis-inhibiting genes (*SOX4* and *HES5*) (Braccioli et al., 2018) in RRMS OPCs, as compared to SPMS OPCs (**Fig. 3F**). These results suggest that OPC maturation and myelination were more active in RRMS brains than SPMS brains.

### Astrocytes showed increased expression of reactive genes and decreased expression of antioxidant genes in RRMS over SPMS

Astrocytes (RRMS 7.6%; SPMS 10.5% of total) encompassed 816 DEGs (**Supplemental Table S3**). RRMS astrocytes showed upregulation of marker genes for reactive astrocytes including *GFAP* and *CD44*, as compared to SPMS (**Fig. 4A, B**), suggesting that RRMS astrocytes were more reactive than SPMS astrocytes. Importantly, RRMS astrocytes expressed fewer genes associated with antioxidant pathways (*SLC7A11, ME1*, and *FTH1*) than SPMS astrocytes (**Fig. 4C**). These results are also consistent with SPMS astrocytes reacting to severe oxidative stress that inhibits OPC maturation (Barateiro et al., 2016; French et al., 2009; Maus et al., 2015; Paintlia et al., 2011; Takase et al., 2018) (**Fig. 3F**). CDR analyses extracted 66 VAGs that included upregulation of *C3*, *SPP1* (encoding osteopontin that induces reactive astrocytes) (Gliem et al., 2015; Ikeshima-Kataoka et al., 2018; Kim et al., 2004; Moon and Shin, 2004)), and *PFN1* (which modulates astrocytic morphology and motility (Schweinhuber et al., 2015)) in RRMS astrocytes (**Supplemental Fig. S2, Table S5 and Table S6**). Collectively, these results implicated functional differences between RRMS (reactive phenotype) vs. SPMS (antioxidative phenotype). Of note, immediate-early genes (*FOS*, *FOSL2, JUN*, and *JUNB*) were found as DEGs in RRMS astrocytes over SPMS astrocytes (**Fig. 4C**), which supported the previous identification of *immediate early astrocytes (ieAstrocytes*) that were identified in studies of experimental autoimmune encephalomyelitis (EAE, an animal model of MS), and which increased in prevalence with disease severity (Groves et al., 2018).

**Figure 4.**
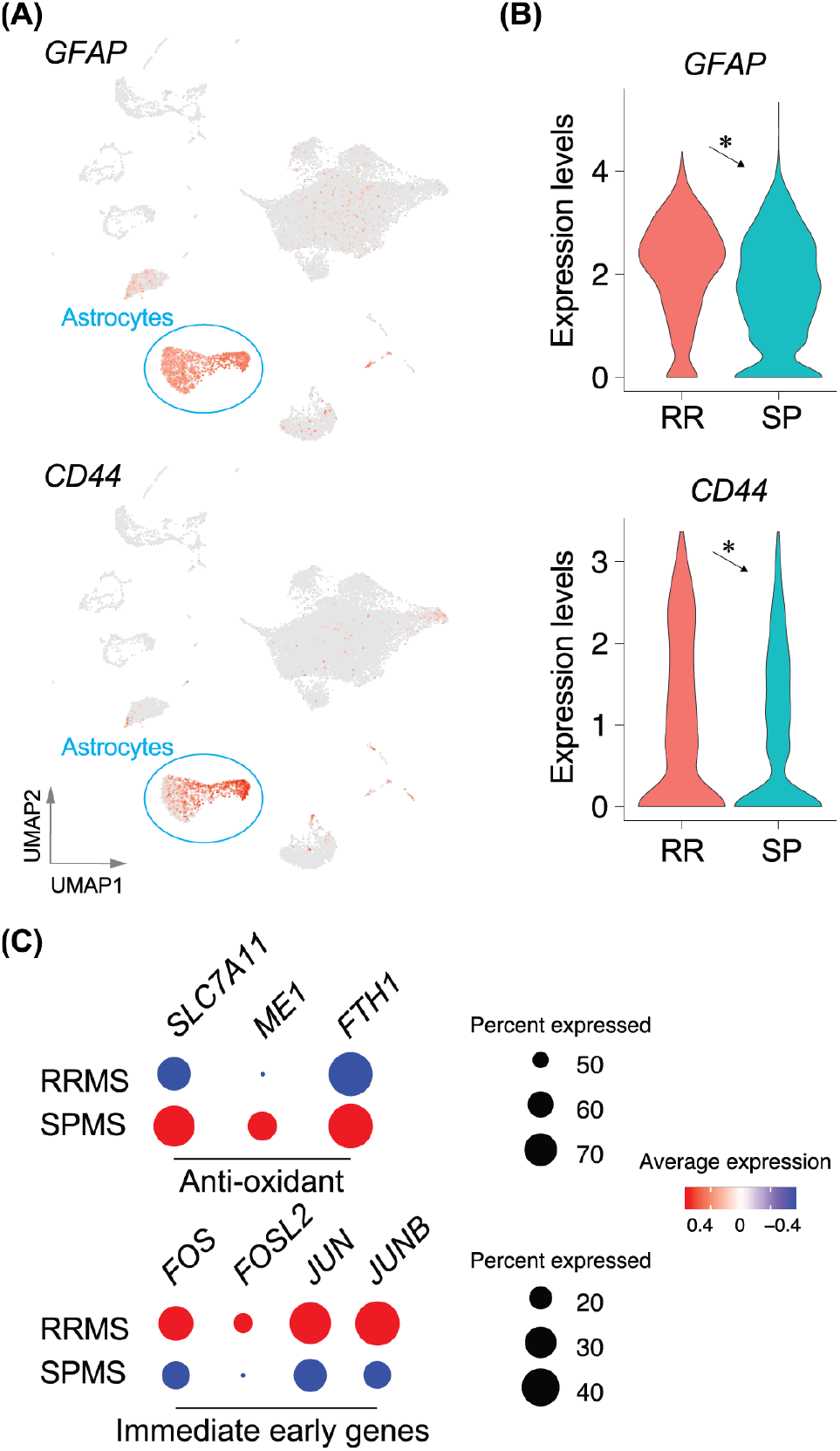
Transcriptomic changes in astrocyte populations between RRMS vs. SPMS. (**A and B**) UMAP (A) and violin plots (B) of astrocyte marker genes. *, adjusted p value < 0.05. (**C**) Dot plots of genes associated with antioxidant pathways and immediate early genes. Size and color indicate the CDRs and expression levels, respectively. Statistical results for DEGs, CDRs and VAGs are provided in **Supplemental Table**.

### Microglia activation genes are increased in RRMS over SPMS

Myeloid cells (RRMS 7.3%; SPMS 10.5% of total) (**Fig. 1E and Supplemental Table S3**), were associated with decreased expression of microglial core genes (*P2RY12, P2RY13*, and *CX3CR1*) in RRMS over SPMS (**Fig. 5A and B**). Activation marker genes for myeloid cells (*CD68, CD74, FTL, APOE*, and *SPP1*) were higher in RRMS than SPMS (**Fig. 5C**), indicating the increased microglial activation in RRMS brains.

**Figure 5.**
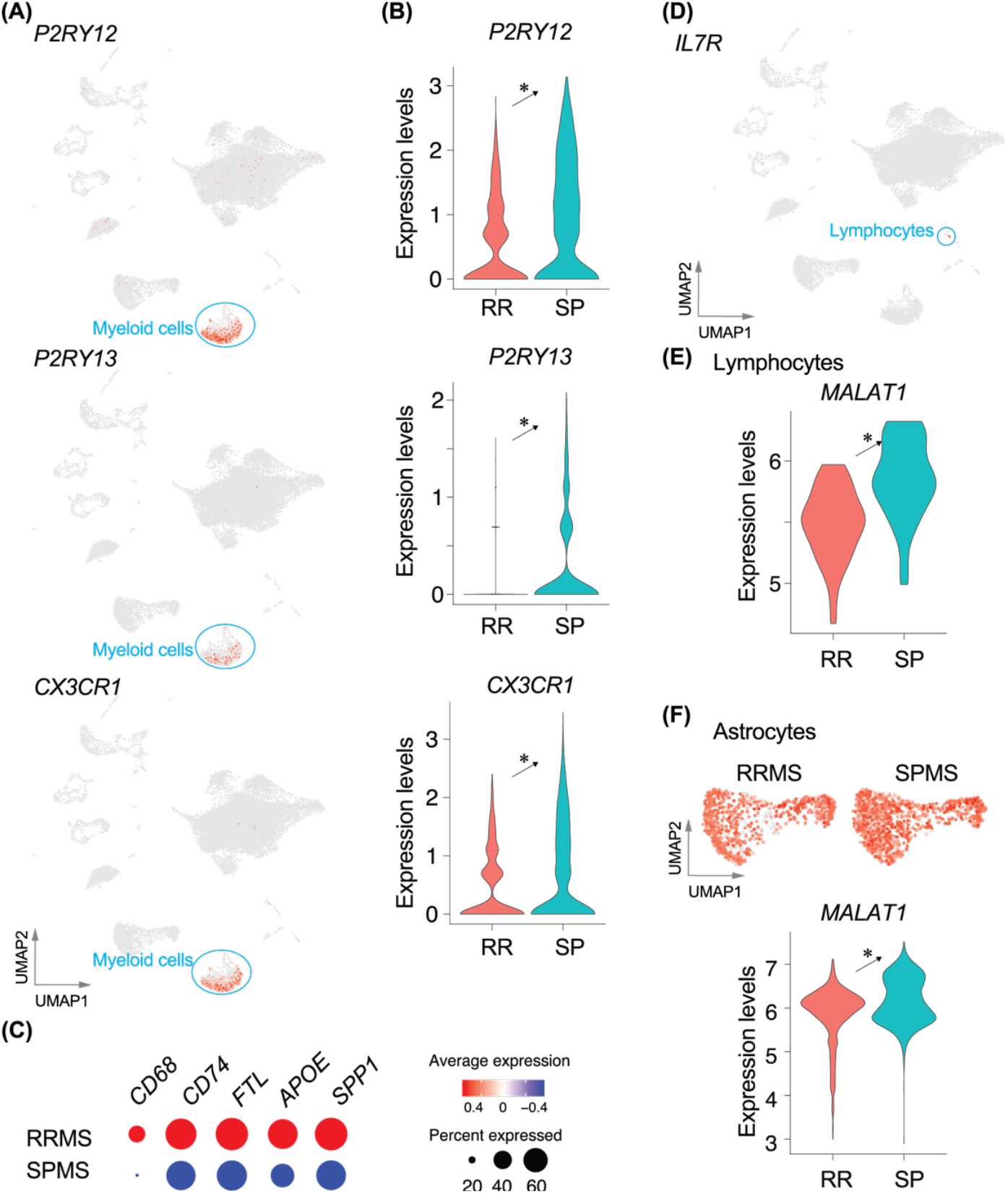
Transcriptomic signatures in other cell types. (**A and B**) UMAP (A) and violin plots (B) of microglial marker genes. *, adjusted p value < 0.05. (**C**) Dot plots of activation marker genes for myeloid cells. Size and color indicate the CDRs and expression levels, respectively. (**D and E**) UMAP of lymphocyte marker gene IL7R (D), and violin plot of *MALAT1* (E). (**F**) UMAP and violin plot of *MALAT1* expression in astrocytes. *, adjusted p value < 0.05. Statistical results for DEGs, CDRs and VAGs are provided in **Supplemental Table**.

### SPMS lymphocytes and astrocytes are possible sources of a SPMS marker gene, MALAT1

Pericytes (~0.5% of total), endothelial cells (~0.6% of total), and lymphocytes (~0.2% of total,) were detected (**Fig. 1E and Supplemental Table S3**), which expressed fewer DEGs (8, 35 and 1 DEGs, respectively) than other cell types. The only DEG found in lymphocytes (**Fig. 5D**) was long non-coding *MALAT1* that was elevated in SPMS (**Fig. 5E**), supporting a previous report identifying *MALAT1* as a potential biomarker for SPMS diagnosis (Shaker et al., 2019). Because the upregulation of *MALAT1* was also found in astrocytes (**Fig. 5F**), both lymphocytes and astrocytes might be possible sources of increased *MALAT1* expression in SPMS.

### A test case for dataset functional validation: reduced sphingosine kinases in SPMS may explain fingolimod’s ineffectiveness as supported by animal knockout studies

Fingolimod is a sphingosine 1-phosphate (S1P) receptor modulator that upon phosphorylation by sphingosine kinases, becomes an active agent (Brinkmann et al., 2010; Chun et al., 2019; Cohen and Chun, 2011; Kappos et al., 2018; Kihara et al., 2015b). It was the first oral agent approved to treat relapsing forms of MS; however, it failed to reach its primary endpoint in Phase 3 clinical trials in primary progressive MS (PPMS) patients (Lublin et al., 2016). In contrast, another S1P receptor modulator, siponimod, successfully obtained approval for treatment of SPMS (Chun et al., 2020; Derfuss et al., 2020). Fingolimod’s lack of efficacy in progressive MS might be explained by alterations in CNS genes that affect fingolimod activity. Two such genes are sphingosine kinases 1 and 2 (*SPHK1* and *SPHK2*) that phosphorylate fingolimod into its active state, and which were compared between RRMS and SPMS brains (**Fig. 6A-E**), revealing reductions in SPMS (~50% for *SPHK1* and ~80% for *SPHK2*) compared to RRMS. Other genes associated with fingolimod metabolism (*SPNS2, S1PR1, S1PR3, S1PR4, S1PR5, SPHK1* and *SGPP1*; **Fig. 6B**) were highly enriched in the top 30 CDR genes of “RRMS > SPMS” groups, markedly contrasting with their detection in “SPMS > RRMS” groups (**Supplemental Fig. S3**).

**Figure 6.**
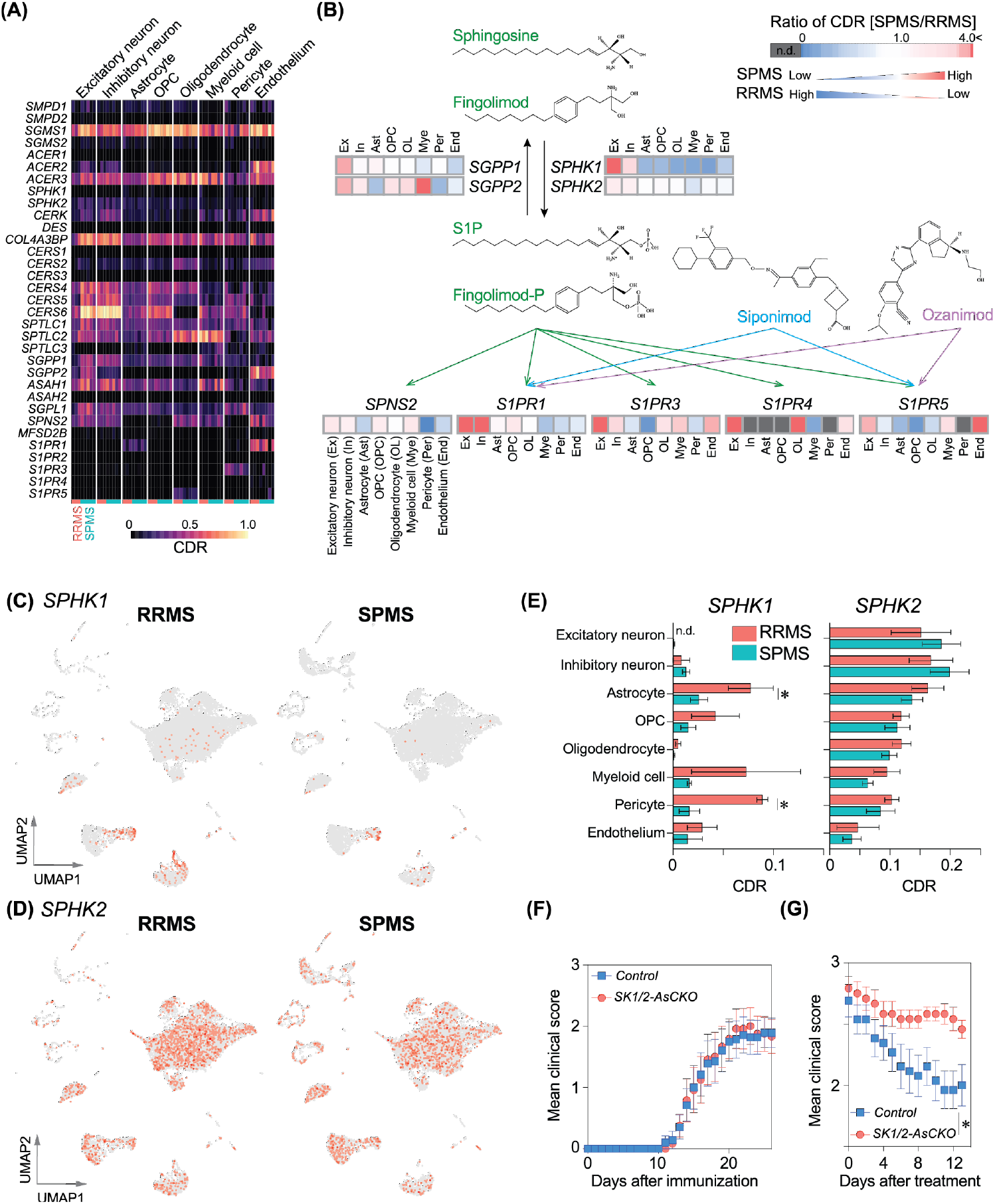
A potential role of CNS-derived sphingosine kinases (*SPHK1/2*) in fingolimod efficacy. (**A**) Heatmap of CDRs for sphingolipid pathway genes. (**B**) Comparison of CDR ratio between SPMS over RRMS in a pathway associated with the mechanism of action of S1P receptor modulators, fingolimod phosphate (fingolimod-P), siponimod, and ozanimod. (**C and D**) UMAP plots of *SPHK1* (C) and *SPHK2* (D). (**E**) CDRs for *SPHK1* and *SPHK2*. n = 3 RRMS vs 5 SPMS. *, p < 0.05 by t-test. Statistical results for DEGs, CDRs and VAGs are provided in **Supplemental Table**. (**F**) EAE disease course in Sphk1/2-AsCKO mice vs. control WT mice. n = 12 and 14 animals, respectively. (**G**) EAE clinical course of fingolimod-treated Sphk1/2-AsCKO mice vs. control WT mice. n = 6 animals. *, p < 0.0001 by two-way ANOVA.

To validate the functional significance of altered astrocyte sphingosine kinase gene expression, astrocyte-specific *Sphk1/2* conditional knockout (SK1/2-AsCKO) mice were challenged with EAE. The disease course in SK1/2-AsCKO mice was equivalent to their wild-type (WT) controls (**Fig. 6F**), although S1P levels in EAE-induced SK1/2-AsCKO spinal cords (3.9 ± 2.1 pmol/mg tissue, n= 3) were significantly lower than those of WT spinal cords (11.3 ± 1.4 pmol/mg tissue, n = 4, *p* < 0.05 by t-test). These results suggested that astrocyte derived S1P was not involved in EAE pathogenesis. In contrast, fingolimod was not effective in EAE mice lacking astrocytic *Sphk1/2* genes (treatment, p < 0.0001; time, p = 0.003; interactions, p = 0.78; by two-way ANOVA, **Fig. 6G**). These results implicated astrocyte SPHK1/2 in fingolimod efficacy and offer an explanation for its lack of efficacy in PPMS.

## Discussion

MS is diagnostically characterized by detectable brain lesions, but global changes away from lesions, including grey matter and volumetric loss, are now accepted as part of MS disease manifestation (Bermel and Bakshi, 2006; Geurts et al., 2012). The use of normal-appearing, region-matched brain samples for snRNA-seq enabled generation of comparable cell clusters between conditions as demonstrated by highly overlapping UMAP layouts between RRMS vs. SPMS (**Fig. 1D**). By contrast, previous snRNA-seq datasets focusing on MS lesions displayed non-overlapping distributions of cells (Jakel et al., 2019; Masuda et al., 2019; Schirmer et al., 2019). Our approach revealed pervasive transcriptomic changes with disease state that were independent of discernible lesion activity.

### Neural vulnerability in MS brains

Prior snRNA-seq studies of MS lesions compared to non-diseased control brains identified selective neuronal loss, severe cellular stress in OLs, increased reactive astrocytes, and microglial activation (Schirmer et al., 2019). These findings were also observed here along with additional features. First, neuronal vulnerability (loss of excitatory neuron marker genes) was found in normal-appearing RRMS brains, suggesting that RRMS brains had already commenced neuropathological changes irrespective of lesions (**Fig. 2**). Second, the SPMS excitatory neurons contrasted with neuronal vulnerability initiated from the RRMS phase. This difference might reflect a more normal state of surviving neurons that were less exposed to RRMS insults, but may still involve axonal degeneration and impairment of neural network rewiring produced by upregulation of ion channels to produce channelopathy-like defects in SPMS (Kumar et al., 2016; Roostaei et al., 2016; Zhong et al., 2016).

### OPC dysfunction in SPMS

Approximately 5% of OPCs were identified from the normal-appearing MS brains, which was comparable to another study (7.8% in MS lesions vs. 6.0% in controls) (Schirmer et al., 2019). Upregulation of *SOX4* and *HES5* in SPMS OPCs, which inhibits OPC differentiation (Braccioli et al., 2018; Munoz-Esquivel et al., 2019; Potzner et al., 2007), along with downregulation of myelin formation-promoting genes, were identified (**Fig. 3**). These results suggest that inhibitors targeting the SOX4-HES5 signaling pathway may be useful for promoting remyelination and delaying MS progression.

### Astrocyte types in MS

A proposed classification of reactive astrocytes sub-divided into neurotoxic A1 astrocytes vs. helpful A2 astrocytes was based on gene expression patterns including immunolabeling for C3^+^, MX1^+^, or CFB^+^ of A1 astrocytes in MS lesions (Liddelow et al., 2017). The universality of this classification is unclear considering that low mRNA expression of A1 markers is found in MS lesions (Schirmer et al., 2019). Moreover, A1 or A2 reactive astrocyte genes were rarely found in DEGs or CDRs in the present study, except for *C3* that was more enriched in microglia rather than astrocytes (**Supplemental Fig. S4**). A1 astrocyte formation has been proposed to require three microglia-derived factors (IL1α, TNF-α, and C1q (Liddelow et al., 2017)) that were not identified in a prior study (Schirmer et al., 2019). However, considerable C1q gene expression (*C1QA*, *C1QB*, and *C1QC*) in microglia, along with minor expression of *IL1A* and *TNF*, was observed here (**Supplemental Fig. S4**). These gene expression differences may reflect technical issues including the 6X increased gene discovery per cell in the current study. Collectively, the A1/A2 classification was only partially supported by some CDRs in A1 gene sets compared to A2 gene sets, indicating a need for further study in assessing MS pathogenesis related to A1/A2 astrocytes.

An alternative and not mutually exclusive astrocyte classification is *ieAstrocytes* (immediate early astrocytes (Groves et al., 2018)) that were defined in EAE spinal cords; *ieAstrocytes* were identified using an unbiased *in vivo* c-Fos activity screen that showed increased *ieAstrocyte* prevalence with worsening disease. *ieAstrocyte* transcriptomic gene expression was supported by the present study (**Fig. 4**) through *FOS* (c-Fos)- expressing astrocytes documented in normal-appearing MS brains, along with upregulation in RRMS over SPMS (**Fig. 4C**). Datasets from prior studies also supported this concept through *FOS* upregulation in astrocytes of MS lesions (Schirmer et al., 2019). The current report indicates that *ieAstrocyte* characteristics extend beyond lesions to normal-appearing regions of the MS brain (**Fig. 4C**).

### Implications for the efficacy of S1P receptor modulators

A notable finding here was the perturbation of sphingolipid metabolic genes, particularly the reduction of *SPHK1/2*-expressing cells in the SPMS brain (**Fig. 6**). This was consistent with a prior study (Schirmer et al., 2019) where *SPHK1/2* expression was lower in chronic inactive lesions (predominately in SPMS (Frischer et al., 2015)) than acute/chronic active lesions (found more often in RRMS (Frischer et al., 2015)). Reduction of *SPHK1/2*-expressing cells in the CNS may, in part, account for the clinical trial failure of fingolimod for PPMS treatment (Lublin et al., 2016). Prior preclinical data showed that the preventative effects of fingolimod in EAE was abolished in constitutive *Sphk2*-deficient mice (Imeri et al., 2016), demonstrating a requirement of SPHK activity for the efficacy of fingolimod in EAE. A direct CNS mechanism of action for fingolimod (Chun et al., 2019; Kihara, 2019) was further supported by functional animal studies demonstrating a loss of fingolimod efficacy in SK1/2-AsCKO mice in a therapeutic paradigm (**Fig. 6G**) which supports the relevance of reduced *SPHK1/2* gene expression in human SPMS brains (**Fig. 6A-E**) to explain progressive MS trial failures. These new data also complement prior reports on fingolimod accumulation in the CNS (Foster et al., 2007) and a requirement for astrocyte S1P_1_ on EAE development (Choi et al., 2011; Groves et al., 2018). In addition, the S1P transporter SPNS2 (spinster 2), which transports not only S1P but also fingolimod phosphate (Hisano et al., 2011), was also significantly reduced in SPMS astrocytes as compared to RRMS astrocytes (**Fig. 6 and Supplemental Table S5**). Thus, fingolimod activity appears to involve local phosphorylation in astrocytes along with related metabolic steps of secretion via SPNS2, and S1P_1_ functional antagonism in an autocrine/paracrine manner. In contrast to fingolimod, next generation S1P receptor modulators do not require phosphorylation by SPHKs for their activity (Chun et al., 2019; Kihara, 2019; Kihara et al., 2014; Kihara et al., 2015b), explaining the efficacy of at least one tested agent, siponimod, in progressive MS (Kappos et al., 2018). The mechanisms underlying the reduction of SPHK-expressing CNS cells in MS brains requires further study but may represent a novel area for therapeutic intervention targeting astrocytes, particularly *ieAstrocytes* (Groves et al., 2018). These results complement an accompanying paper with an exploration of astrocytic S1P_1_ signaling (Jonnalagadda, 2021), which identified a linkage between the sphingolipid and vitamin B_12_ metabolisms. Both papers strongly support that astrocytes play central roles for fingolimod’s non-immunological mechanism of action.

Taken together, the current snRNA-seq dataset complements prior snRNA-seq MS lesion datasets to reveal neural vulnerabilities within normal-appearing MS-affected brains. The perturbations of sphingolipid signaling pathways may help to explain the lack of efficacy for fingolimod in PPMS and implicates non-prodrug strategies to impact sphingolipid signaling in progressive CNS disorders.

## Acknowledgments

We thank Dr. Shaun R Coughlin for providing Sphk1/2^flox/flox^ mice and Dr. G. Kaeser and D. Jones for editing the manuscript. This work was supported by National Institute of Neurological Disorders and Stroke of the National Institutes of Health (NIH) under award numbers R01NS084398 (J.C.), R01NS103940 (Y.K), R01NS096148 (R.D.), and R35NS097303 (B.D.T); the National Institute of Mental Health of the NIH under award number R01MH051699 (J.C.); the Department of Defense under award number W81XWH-17-1-0455 (J.C.); and research grants from Novartis (J.C.). The content is solely the responsibility of the authors and does not necessarily represent the official views of the National Institutes of Health.

## Author Contribution

Y.K., Y.Z., D.J., W.R., C.P., B.S., and R.R. performed experiments, analyzed data, and wrote the paper. R.D. and B.D.T. collected human MS brains. J.C. conceived of the project and wrote the paper with the co-authors. All authors contributed towards the final version of the manuscript.

## Materials and Methods

### Human brain tissues

Postmortem prefrontal cortices (BA10) from RRMS and SPMS donors were collected, frozen immediately, and stored at −80°C at the Cleveland Clinic and the guidelines of the Cleveland Clinic human research ethics committee were followed (Dutta et al., 2019). The detailed sample information is provided in **Supplemental Table S1**.

### Nuclear isolation, 10x Genomics platform, and short-read sequencing

Tissues were cut on a cryostat (Leica) and a total of 300 μm sections from each were stored at −80 °C until use. The buffers used in this procedure were made in autoclaved diethyl pyrocarbonate (DEPC; MilliporeSigma, Cat # D5758)-treated water. All the steps from here on were performed at 4 °C with additional equilibration time at each step. The samples were initially placed in 0.5 mL nuclear extraction buffer (NEB; 0.32 M Sucrose, 5 mM CaCl_2_.2H_2_0, 3 mM (CH_3_COO)_2_Mg.4H_2_O, 0.1 mM EDTA, 10 mM Tris/HCl (pH 8.0), 0.1% Triton X-100 with cOmplete™ mini protease inhibitor cocktail tablet (MilliporeSigma, Cat # 11836153001), and fresh 0.2% RNase inhibitor (Takara Bio, Cat# 2313A)) for 15 minutes. After adding another 0.5 mL NEB, the samples were homogenized thoroughly with a Dounce homogenizer. Upon transferring this homogenate to a new tube, the Dounce homogenizer was rinsed out with 1 mL of NEB. Homogenates were then passed through a CellTrics® 50 μm filer (Sysmex, Cat # 04-004-2324). After washing the filter with NEB once, the filtrates were centrifuged at 1600 x *g* for 5 mins. The resulting pellets were resuspended with 1 mL of PBSE (1X PBS, 2 mM EGTA and fresh 0.2% RNase inhibitor), transferred into a new tube, diluted with PBSE, and centrifuged again at 1600 x *g* for 5 mins. Meanwhile, density gradients of OptiPrep™ (MilliporeSigma, Cat # D1556) were made in FACS tubes with 0.4 mL 35% OptiPrep™ on the bottom and 2 mL 10% OptiPrep™ on the top. Solutions used for density gradients were as follows: OptiPrep™, solution B (6X concentrations: 120 mM Tricine/NaOH pH 7.8, 150 mM KCl and 30 mM MgCl_2_.6H_2_O) and solution D (0.25 M Sucrose in 1X solution B with freshly added 0.2% RNase inhibitor). The OptiPrep™ solutions were diluted with solution D to make the final concentrations to 35% and 10%. Nuclear pellets were resuspended with 0.2 mL solution D, followed by addition of 0.2 mL of 10% OptiPrep™ solutions, which were layered onto the 10% OptiPrep™ solution, and centrifuged at 3250 x *g* for 10 mins. The nuclei that settled at the boundary of 35% and 10% were collected, washed twice with PBSE containing 1% fatty acid-free bovine serum albumin (BSA; Gemini Bio, Cat # 700-107P), passed through a CellTrics® 50 μm filer, washed again with PBSE + BSA, and labeled with DAPI (1:100,000, 4’,6-diamidino-2-phenylindole; MilliporeSigma, Cat # D9542). About 3 × 10^5^ nuclei were sorted by a BD FACSAria™ Fusion (BD Biosciences) into a collection tube that was precoated and filled with 1 mL of PBSE + BSA. The nuclei were washed once and resuspended with PBSE + BSA to obtain around 1500 cells/μL, and the counts were confirmed by Countess™ II (ThermoFisher Scientific). Single-nucleus capture (target capture of 5,000 nuclei per sample) and library preparation was conducted using Chromium Next GEM Single Cell 3’ GEM Library & Gel Bead Kit v3 (10x Genomics, Cat # PN-1000075) and Chromium Single Cell B Chip Kit (10x Genomics, Cat # PN-1000074) according to the manufacturer’s instructions without modification. Singlenucleus libraries were sequenced on the Illumina HiSeq3000 machine at GENEWIZ, Inc. (U.S.),

### Bioinformatics analyses

Raw FASTQ files were input into the Cell Ranger count pipeline (Cell Ranger V3.0.2, 10x Genomics) to align reads to the GRCh38 human genome, quality filter cellular barcodes and unique molecular identifiers (UMIs), and count UMIs by gene. To account for unspliced pre-mRNA transcripts in the nucleus, a custom pre-mRNA annotation file was generated and supplied to Cell Ranger as per 10x Genomics instructions. Gene-UMI count matrices generated from individual samples were filtered to exclude genes detected in less than three nuclei and to retain nuclei with at least 300 genes expressed. Next, dying cells and multiplets were removed by excluding nuclei with > 1% mitochondrial count fraction or transcript number exceeding the 75^th^ quantile + 1.5*IQR (Interquartile Range). After these stringent filtering steps, we obtained a total of 33,197 high quality nuclei from eight MS brains (3 RRMS vs. 5 SPMS). After this initial quality filtering, UMI raw count matrices from individual samples were normalized using a regularized negative binomial regression method (Hafemeister and Satija, 2019) to regress out mitochondrial count fractions and then were integrated into one combined dataset using Seurat V3.0 (Stuart et al., 2019) in R. Next, nuclei in the integrated dataset were clustered through a shared nearest neighbor (SNN) algorithm and visualized in two-dimensional space using uniform manifold approximation and projection (UMAP) (Becht et al., 2018). Cell types were annotated by performing cell label transfer in Seurat based on a reference normal brain snRNA-seq dataset that we previously published (Lake et al., 2018), which produced cell type-specific markers unbiasedly (**Supplemental Fig. S1 and Table S3**). DEGs (differentially expressed genes; FDR-adjusted P < 0.05, log2(fold change) > 1.1) in each cluster were identified by fitting a hurdle model adjusting for sex, age, and RIN scores in MAST (Finak et al., 2015). CDRs (cell distribution rates) were calculated from a transformed UMI count matrix that has undergone a binary transformation based on presence-or-absence of gene expression. The statistical difference of CDR between RRMS and SPMS groups was determined by the nonparametric Wilcoxon rank-sum test. The VAGs (vastly altered genes) were defined as the intersection between DEGs and genes with significant CDRs. Pathway analyses were executed in the Reactome Pathway Database https://reactome.org/ and the results are provided in **Supplemental Tables S7-9**. Fastaq files have been deposited into the Gene Expression Omnibus (GEO, https://www.ncbi.nlm.nih.gov/geo/). The accession number is GSE179590.

### Animal studies

Animal protocols were approved by the Institutional Animal Care and Use Committee of the Sanford Burnham Prebys Medical Discovery Institute and conformed to National Institutes of Health guidelines and public law. The *Sphk1/2*^flox/flox^ mice (Pappu et al., 2007) were crossed with human GFAP-Cre (FVB-Tg(GFAP-cre)25Mes/J, Cat# JAX:004600, RRID:IMSR_JAX:004600) mice to generate astrocyte specific *Sphk1/2* conditional KO (SK1/2-AsCKO: Sphk1/2^flox/flox^:GFAP-Cre) mice. EAE was induced in 8- to 12-wk-old female mice as previously described (Kihara et al., 2015a; Kihara et al., 2009). Briefly, mice were immunized with 100 μL emulsions containing 150 μg myelin oligodendrocyte glycoprotein 35–55 (MOG35–55) (MEVGWYRSPFSRVVHLYRNGK, EZBiolab) in PBS and a mixture of Difco™ incomplete Freund’s adjuvant (BD Biosciences, Cat # 263910) and 4 mg/mL Difco™ Adjuvant (*Mycobacterium Tuberculosis* H37; BD Biosciences, Cat # 231141). Daily clinical scores were given as follows: 0, no sign; 0.5, mild loss of tail tone; 1.0, complete loss of tail tone; 1.5, mildly impaired righting reflex; 2.0, abnormal gait and/or impaired righting reflex; 2.5, hindlimb paresis; 3.0, hindlimb paralysis; 3.5, hindlimb paralysis with hind body paresis; 4.0, hind- and forelimb paralysis; and 4.5, death or severity necessitating euthanasia. Fingolimod was administered via gavage (1 mg/kg; gifted from Novartis). S1P levels were quantified at the Center for Metabolomics and Mass Spectrometry at The Scripps Research Institute, USA.

## Supplemental Figures provided as Supplemental_Figures.pdf

- Supplemental Figure S1. Heatmap of cell type-specific gene expression.
- Supplemental Figure S2. Comparison between DEGs vs. CDR genes in each cell type.
- Supplemental Figure S3. Top 30 CDR genes in the sphingolipid pathway.
- Supplemental Figure S4. Pan/A1/A2-specific genes.

## Supplemental Tables provided as Supplemental_Tables.xlsx

- Supplemental Table S1: Sample information
- Supplemental Table S2: Stats for snRNA-seq results
- Supplemental Table S3: Proportion of cell types and their marker genes
- Supplemental Table S4: DEGs in each cell type
- Supplemental Table S5: Genes identified by CDR analyses and VAGs in each cell type
- Supplemental Table S6: VAGs in each cell type
- Supplemental Table S7: Reactome pathway analysis of DEGs found in excitatory neurons
- Supplemental Table S8: Reactome pathway analysis of VAGs found in OLs
- Supplemental Table S9: Reactome pathway analysis of VAGs found in OPCs

