## Supplemental Figures for "Single-nucleus RNA-seq of normal-appearing brain regions in relapsing-remitting vs. secondary progressive multiple sclerosis"

**Supplemental Figures provided as Supplemental_Figures.pdf**

- Supplemental Figure S1. Heatmap of cell type-specific gene expression.
- Supplemental Figure S2. Comparison between DEGs vs. CDR genes in each cell type.
- Supplemental Figure S3. Top 30 CDR genes in the sphingolipid pathway.
- Supplemental Figure S4. Pan/A1/A2-specific genes.

**Supplemental Tables provided as Supplemental_Tables.xlsx**

- Supplemental Table S1: Sample information
- Supplemental Table S2: Stats for snRNA-seq results
- Supplemental Table S3: Proportion of cell types and their marker genes
- Supplemental Table S4: DEGs in each cell type
- Supplemental Table S5: Genes identified by CDR analyses and VAGs in each cell type
- Supplemental Table S6: VAGs in each cell type
- Supplemental Table S7: Reactome pathway analysis of DEGs found in excitatory neurons
- Supplemental Table S8: Reactome pathway analysis of VAGs found in OLs
- Supplemental Table S9: Reactome pathway analysis of VAGs found in OPCs

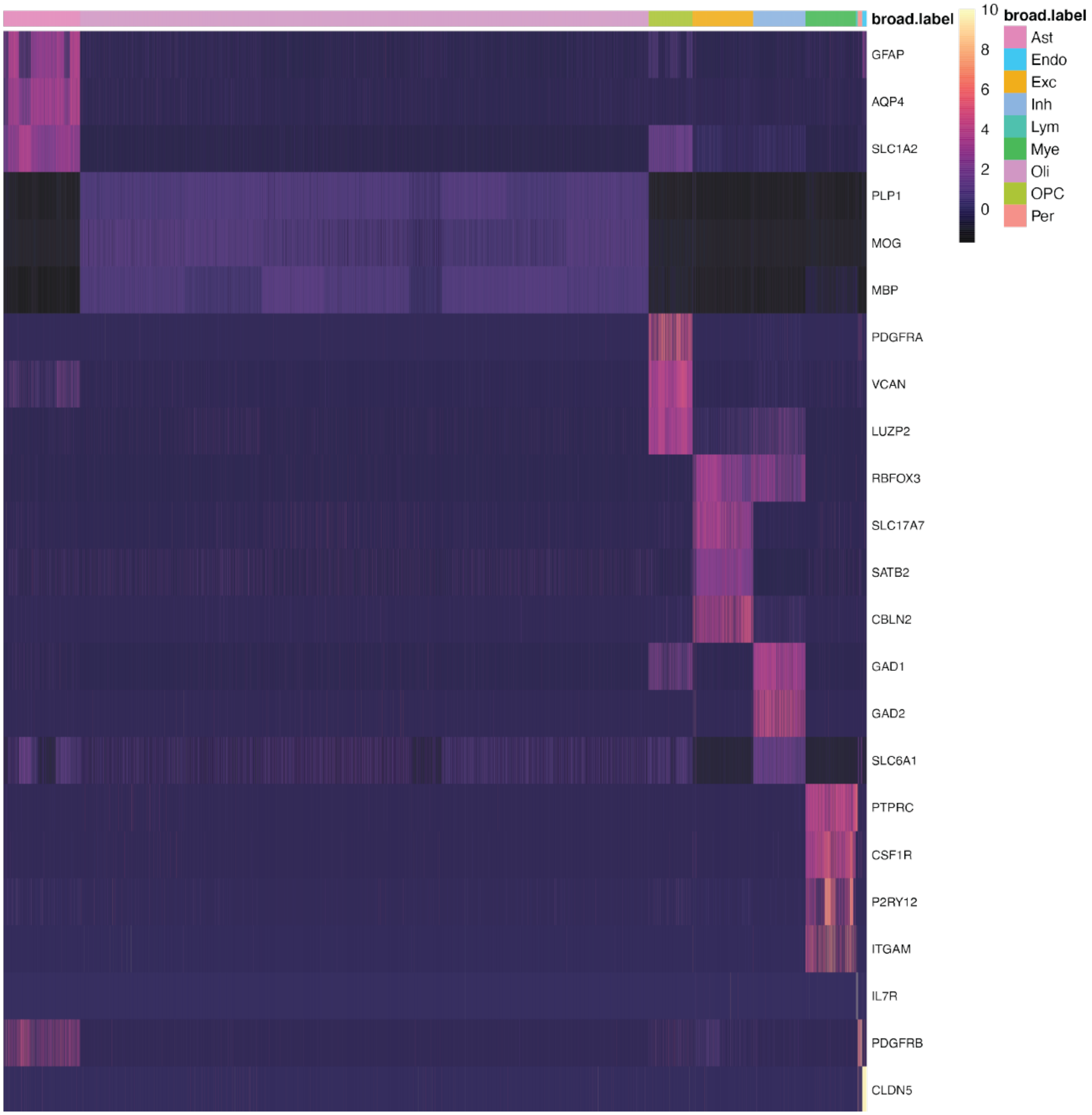


**Figure S1. A Heatmap of cell type-specific gene expression.**

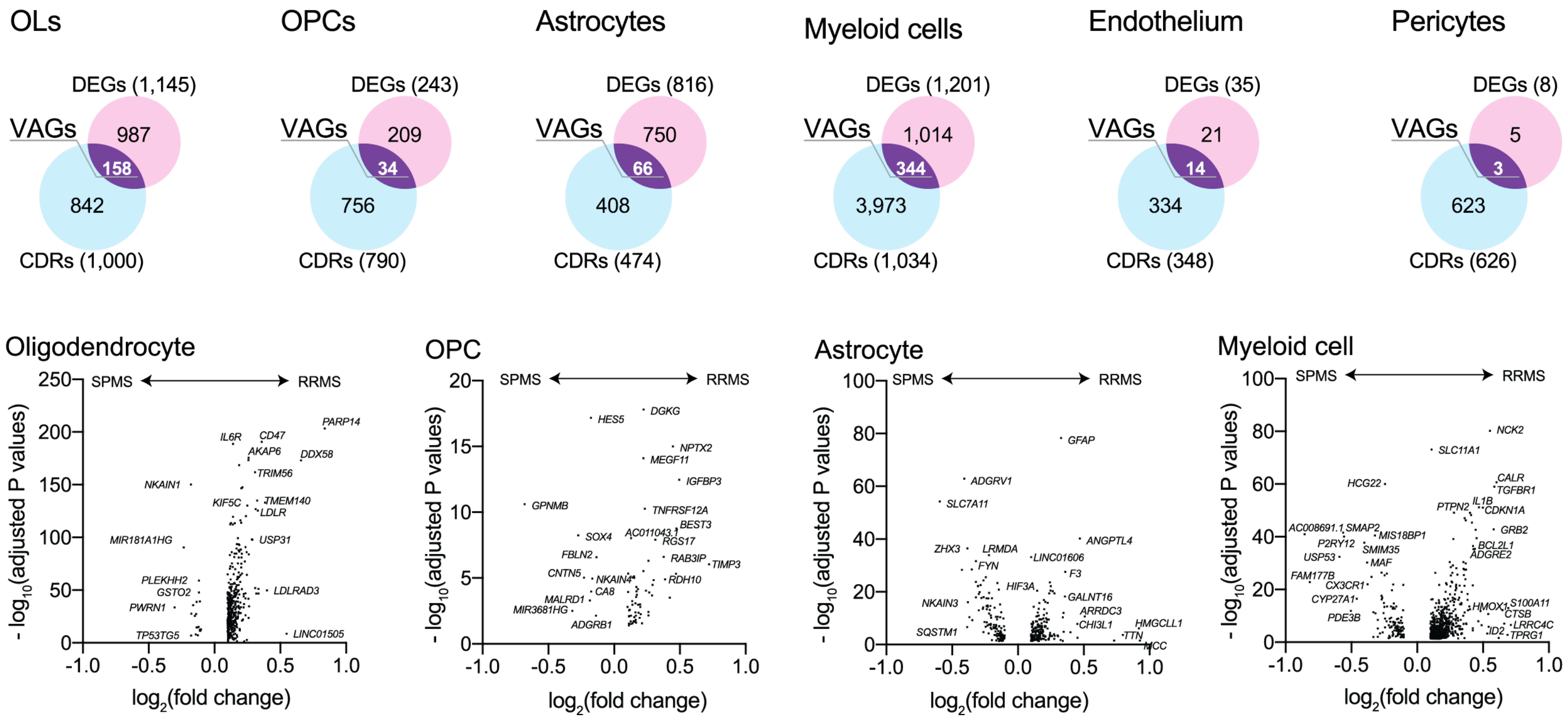


**Figure S2. Comparison between DEGs vs. CDR genes in each cell type.** (Top) Venn diagrams between DEGs and CDRs. The intersections are VAGs. (Bottom) Plots for VAGs in oligodendrocytes, OPCs, astrocytes, and myeloid cells.

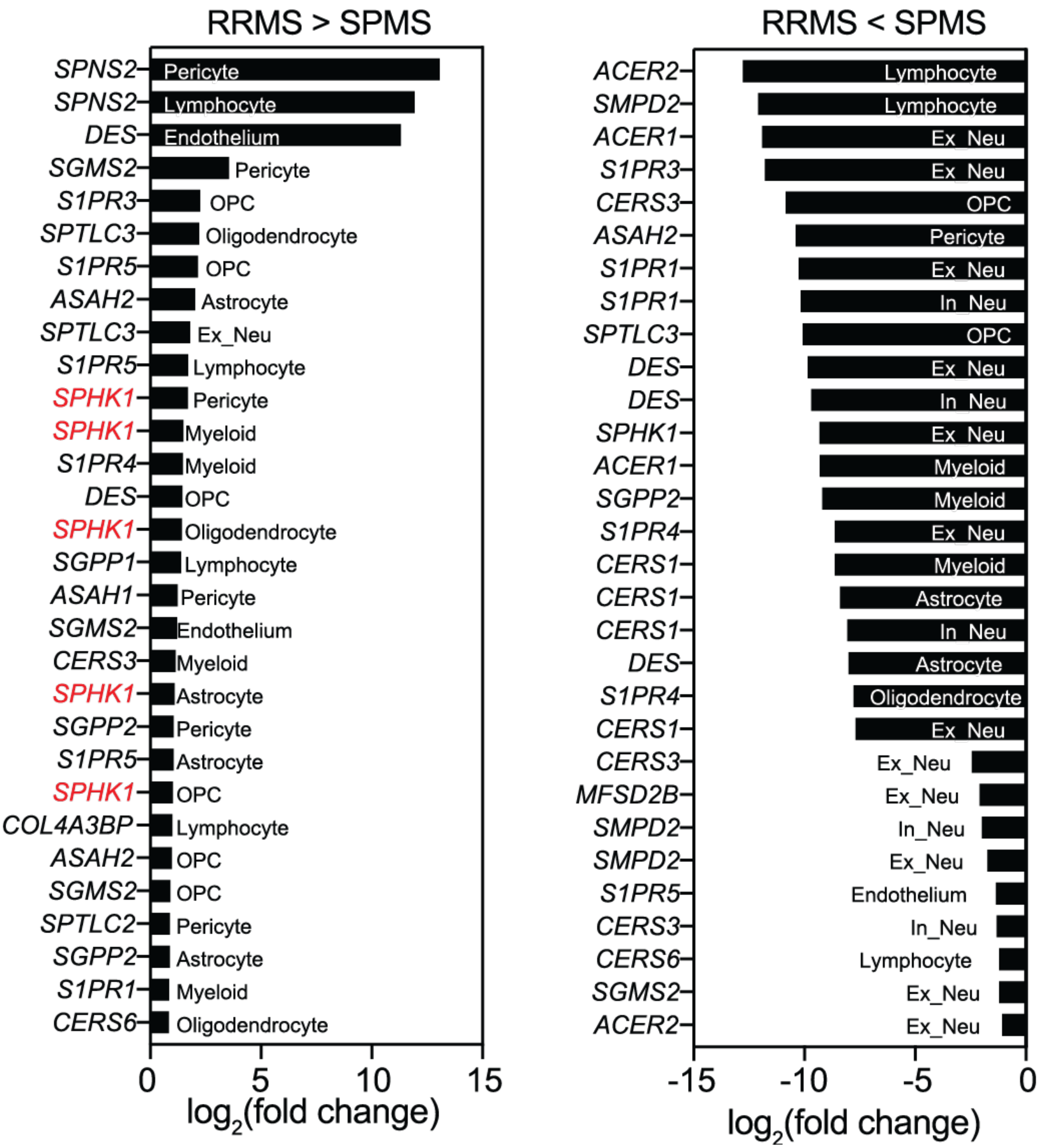


**Figure S3. Top 30 CDR genes in the sphingolipid pathway.**

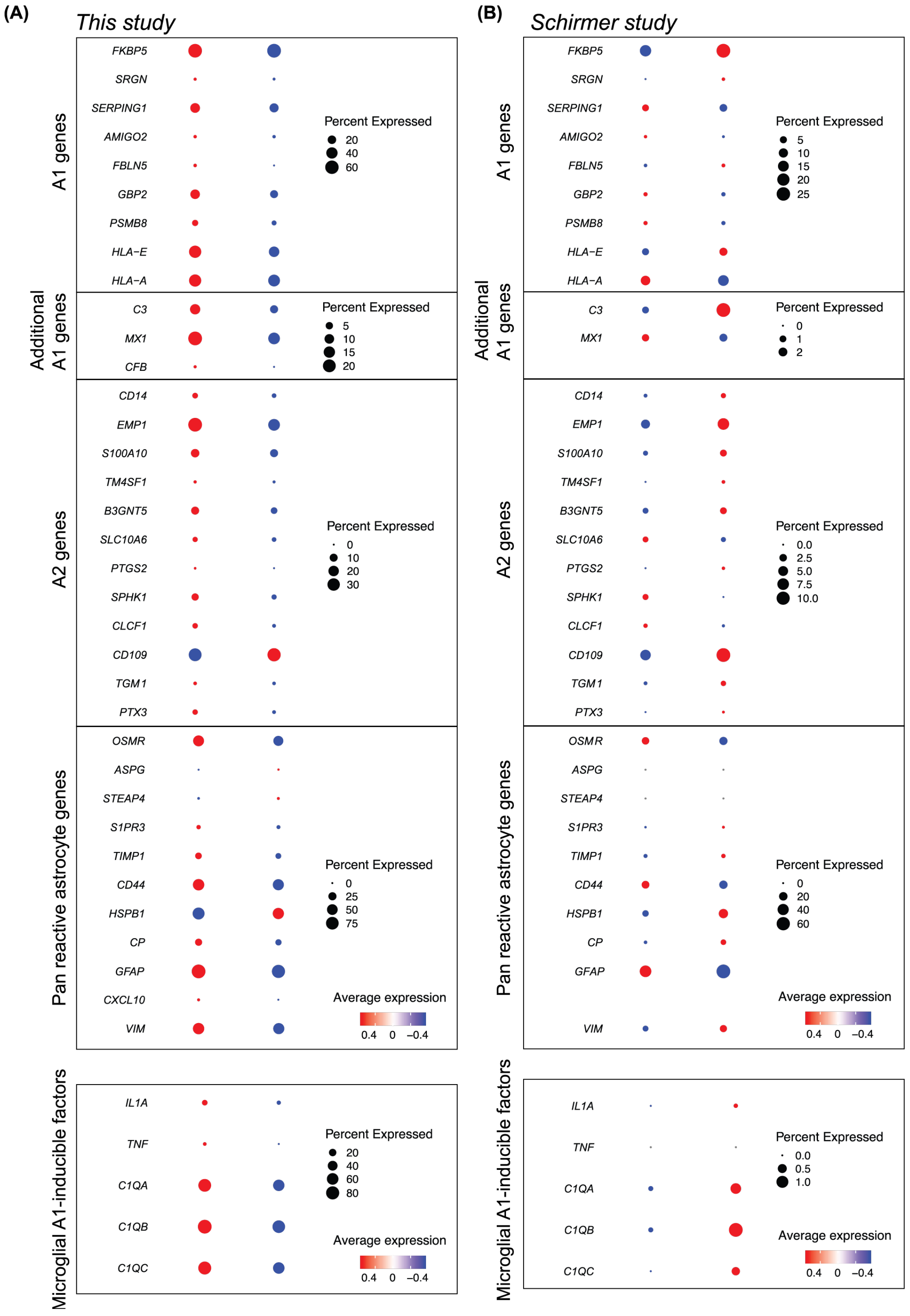


**Figure S4. Astrocytic gene expression associated with A1/A2/Pan reactive astrocytes and expression of microglial A1-inducible factors.**
